# Isolation and Characterization of Salt Tolerant Soil Bacteria from Selected Locations in the West Coast of Sri Lanka

**DOI:** 10.1101/2025.05.01.651645

**Authors:** G.L.C.N.T. Senarath, R. Edirisinghe, M.D. Senarath-Yapa, P.A.I.U. Hemachandra, Y Somaratne, N.U Jayawardana

## Abstract

**Background:** Soil salinity is one of the inherent problems in coastal soils. The increased soil salinity in costal soils results in low bacterial diversity. However, salt tolerant bacteria have the ability to survive in saline soils because of their unique salt tolerant mechanisms. The current study was focused on identifying the salinity tolerant bacteria in the West coast of Sri Lanka. The molecular characterization of potential salt tolerant bacteria was carried out by 16S rRNA sequencing.

**Results:** Negombo lagoon, Balapitiya coastal area and Beruwala coastal area were selected as three locations from the West coast of Sri Lanka. Twenty bacterial isolates namely NE01 to NE09 (Negombo lagoon), BA01 to BA07 (Balapitiya coastal area), BE01 to BE04 (Beruwala coastal area) were obtained from 12 soil samples of selected locations. The isolates were characterized based on the bacterial colony morphology, Gram’s stain and biochemical tests namely, catalase test, modified oxidase test, hemolytic reaction on blood agar and growth on MacConkey agar. All the bacterial isolates were screened for salt tolerance at different concentrations of NaCl ranging from 3 dS/m to 18 dS/m. All 20 bacterial isolates were found salt tolerant upto the highest NaCl concentration (18 dS/m). The salinity tolerant behavior of selected five isolates namely NE 03, NE 06, NE 07, BA 01 and BA 02 were observed by enumerating the number of colony forming units per milliliter (CFU/ml) with increasing salinity. Molecular identification of the above five isolates were carried out by sequencing the 16S rRNA gene and were identified upto their genus level as *Bacillus sp*., *Aeromonas sp*., *Pseudomonas sp*., *Aeromonas sp.* and *Mangrovibacter* sp. respectively.

**Conclusions:** The identified salt tolerant bacterial species can be screened for their plant growth promoting rhizobacterial (PGPR) activities such as phosphate solubilization, potassium solubilization, indole acetic acid (IAA) and nitrogen fixation. Therefore, there is a potential for using these isolates in biofertilizer formulations with the aim of increasing the productivity of saline affected soils of Sri Lanka.

## Background

Soil salinity refers to the accumulation of water-soluble salts in soils. Examples of responsible ions for the formation of salts that accumulate in soil are potassium (K^+^), calcium (Ca^2+^), magnesium (Mg^2+^), sodium (Na^+^), sulfate (SO_4_^2-^), chloride (Cl^-^), carbonate (CO_3_^2-^), and bicarbonate (HCO_3_^-^) (Orhan and Gulluce, 2015). Soil salinity is an enormous problem to coastal regions because it impacts the agricultural production, biodiversity and sustainable development (Gopalakrishnan and Kumar, 2020).

As Sri Lanka is an island, it has a coastline of 1562 km, and it could be more than 2000 km if the coastlines of lagoons, bays and inlets are added (Koralagama,2008). The coastline of Sri Lanka consists of a wide range of geomorphological features such as headlands, bays, lagoons, peninsulas, spits, bars and islets (Lowry and Wickremeratne, 2012). When considering the arable land extent in Sri Lanka, approximately 11,200 ha of coastal lands have been affected and more than 50% of the coastal paddy lands have been abandoned due to salinity (Perera *et al*., 2018). The study is focused on the West coast of Sri Lanka which has highly productive coastal ecosystems such as lagoons, estuaries and mangroves (Perera *et al*., 2018). These coastal landscapes also become victims of coastal salinity. The saline soil is defined as the soil having electrical conductivity (EC) of the saturation soil extract (EC_e_) more than 4 dSm^-1^ at 25 °C and exchangeable Na 15% (Perera *et al*., 2018), The electrical conductivity (EC) of the saturation soil extraction in the west coast of Sri Lanka also has exceeded 4 dS/m (Perera *et al*., 2018).

In order to enhance the productivity of these coastal agricultural fields, there are specific bacterial groups that are capable of surviving and aid in the recycling of nutrients (Orhan and Gulluce, 2015). This specific group of bacteria is called salt tolerant bacteria. They utilize a wide range of carbon sources that are suitable for alkaline environments (Yu *et al*., 2022).Based on the NaCl concentration that they can tolerate, salt tolerant bacteria are categorized into 4 groups as non-halophilic, slight halophilic, moderate halophilic and extreme halophilic (Kumawat *et al*., 2022). Salt tolerant bacterial species isolated so far from saline coastal soils belong to the genera *Alcaligenes* (Sharma *et al*., 2021)*, Pseudomonas* (Kumawat *et al*., 2022)*, Azospirillum* (Rath *et al*., 2019)*, Bacillus* (Kumawat *et al*., 2022) and *Klebsiella* (Orhan and Gulluce, 2015).

The salt tolerant bacteria offer numerous actual or potential applications in various fields of biotechnology, as a result of their adaptation to saline environments (Margesin and Schinner, 2001). At present, salt tolerant bacteria can be used to resist environmental stress due to high salt concentration. They also play an important role in the treatment of high salt sewage, developing bioactive substances and salt tolerant enzymes for biological application (Margesin and Sehinner,2001; Yu *et* al., 2021).

According to the study Sharma (2021), the effects of salt stress on plants can be minimized with the use of potential salt tolerant plant growth promoting rhizobacteria (PGPR). The PGPR are heterogenous bacteria which encourage plant growth by numerous mechanisms such as phosphate solubilization, nitrogen fixation, production of indole acetic acid (IAA) and phytochrome production (Orhan and Gulluce,2015; Kumawat *et* al., 2022). These bacteria act as biofertilizers which can be utilized to obtain a better crop yield in salt affected regions specifically in coastal agricultural lands. (Silva *et al*., 2013)

In addition to that, plants can be engineered with salt tolerant genes isolated from salt tolerant bacteria to boost the crop productivity in salt affected coastal agricultural lands (Agarwal *et al*., 2013).

## Methods

### Collection of soil samples

A total of 12 soil samples (four samples per site) were collected from three selected locations on the West Coast of Sri Lanka namely Negombo (Location 01, 7.2064863’ North, 79. 8356403’ East), Balapitiya (Location 02, 6.1641008’ North, 80.0230447’ East) and Beruwala (Location 03, 6.2644584’ North, 79.5926793’East) [Fig. 1]. The sampling sites were within 20 meters from the shore. The distance between two soil sampling spots was three meters. The sampling depth was 10 cm and the weight of one soil sample was 200 grams. The soil samples were collected into sterile ziplock bags and stored inside a cooling container during transportation. The samples were stored at 4 ° C until analysis.

**Fig 1:**
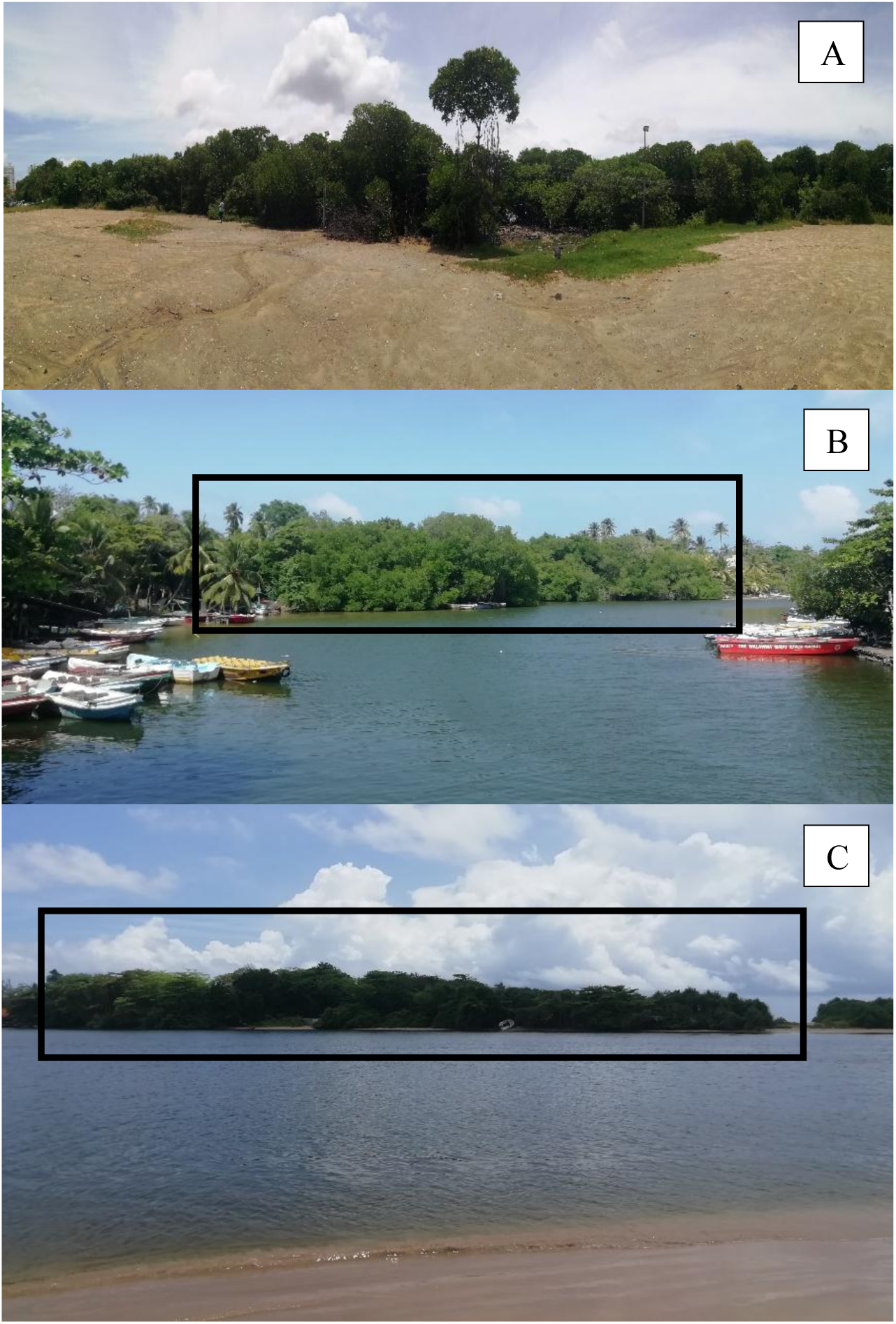
Selected soil sampling locations in the West Coast of Sri Lanka; Negombo (A), Balapitiya (B), Beruwala (C).

### Analysis of soil samples for pH and electrical conductivity

The pH and electrical conductivity (EC) of soil solution: water ratios of 1:1 were determined using the standard method described by ‘Jackson, M. L. (1973). *Soil chemical analysis*.

### Isolation of bacteria from soil samples

Bacteria were isolated from the soil samples on nutrient agar (NA) medium using a serial dilution followed by the spread plate technique. Dilutions up to 10^-3^ were plated and incubated for 24 h at 30 °C. Single isolated colonies were selected for subculturing and their macromorphological colony characters were observed (Table 2). Pure cultures of bacteria were obtained by repeated subculturing.

**Table 1:** Classes of salinity and EC in dS/m (adapted from NRSC soil survey handbook: https://www.nrcs.usda.gov/resources/guides-and-instructions/national-soil-survey-handbook).

| Saturate extract (dS/m) | Salinity class |
| --- | --- |
| < 2.0 | Non-saline |
| 2-4 | Slightly saline |
| 4-8 | Moderately saline |
| 8-16 | Strongly saline |
| >16 | Very strongly saline |

**Table 2:**
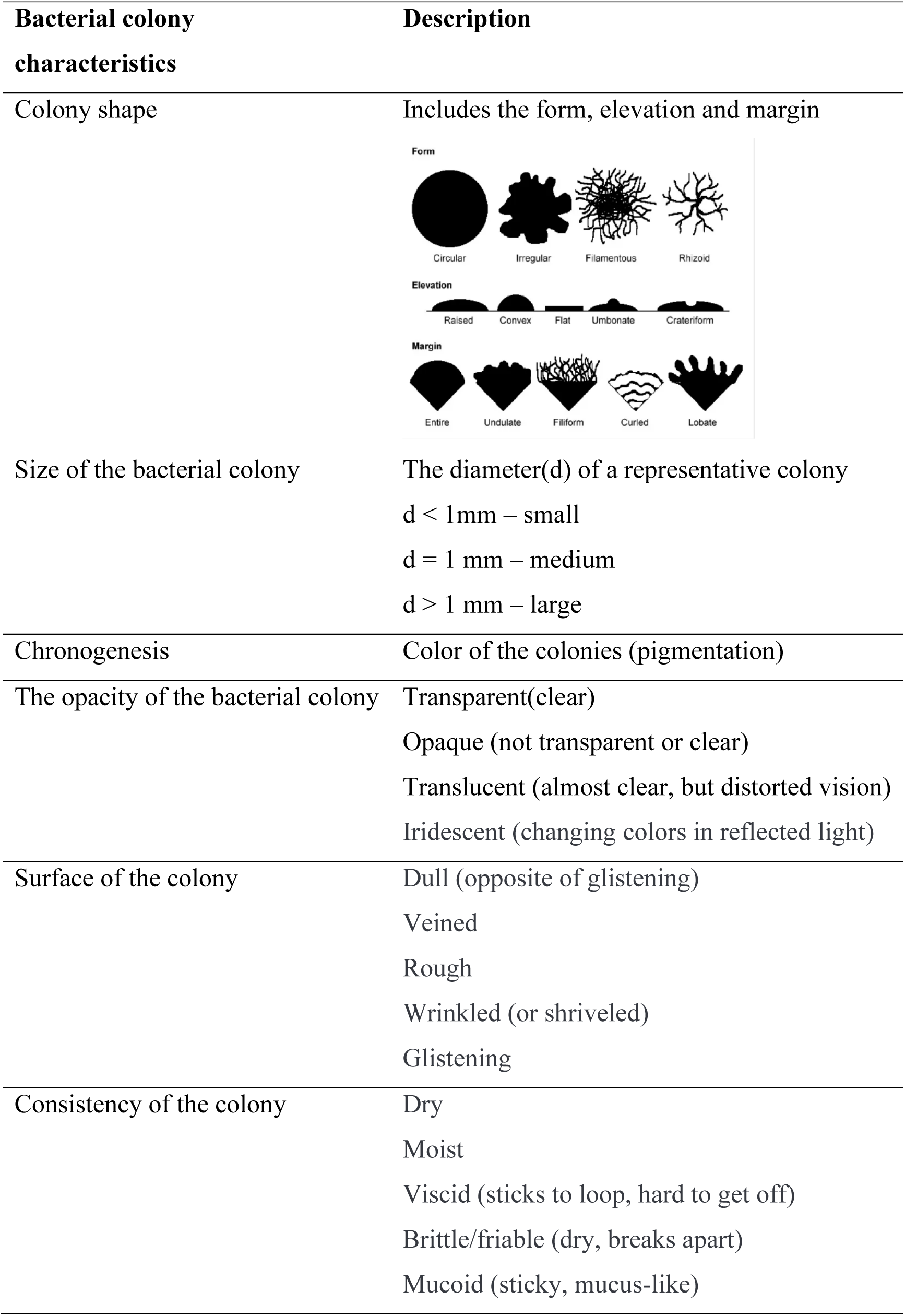
Macromorphological colony characters of bacterial isolates (source: https://www.researchgate.net/figure/Morphological-characteristics-of-different-bacterial-isolates_tbl1_277674878).

### Characterization of bacterial isolates

Gram’s stain test was performed to determine the Gram status of the isolates. Additionally, Catalase test, modified oxidase test, hemolytic reaction on blood agar, and growth on MacConkey agar (only for gram-negative isolates) were used for further characterization of the isolates.

### Screening of bacteria for salt tolerance

The isolates were screened for salt-tolerance by spot inoculating the isolates on Luria Bertani Broth (LB broth) containing different concentrations of sodium chloride (NaCl). Four salinity levels [(slightly saline (3 dS/m), moderately saline (6 dS/m), strongly saline (12 dS/m), and very strongly saline (18 dS/m)] were prepared by adding calculated amounts of NaCl to LB broth. Isolates were spot inoculated and incubated at 25 °C ± 2 °C for 5 days. After 5 days of incubation, each solution was cultured on nutrient agar by using streak plate technique and observed for growth.

### Measuring the growth of isolates along with salinity

After 5 days of incubation, the isolates that are salt tolerant were selected based on absorbance values at 600 nm (Optical Density at 600 nm). LB broth with the highest EC value (18 dS/m) was taken as the growth medium for the selected isolates to measure and compare their growth differences under saline conditions. After inoculating an isolate into LB solution which has the EC value of 18 dS/m, initial plate count was taken as Colony Forming Units (CFU) per milliliter. Then the inoculated broth was incubated at room temperature (25 ℃ ± 2 ℃) for five days and after the incubation period, the final plate count was taken. The salt tolerance of the isolates was compared by the difference between the initial and final CFU per milliliter. CFU/ml was obtained by the following equation, CFU/ml = [(average number of colonies) * (dilution factor)] / (volume of the inoculum in ml).

### Molecular characterization of bacterial isolates

Isolation of genomic DNA: Bacterial genomic DNA was isolated from 24-hours old broth cultures (10 ml) of bacterial isolates using HiPurA^TM^ Bacterial Genomic DNA Purification Kit (MB505-5prp).

### Amplification and sequencing of 16s rRNA

Universal primers: forward primer, 27F (5’ AGA GTT TGA TCC TGG CTC AG 3’, 25 nmole DNA oligo, 20 bases) and reverse primer, 1492R (5’ GGT TAC CTT GTT ACG ACT T 3’, 25 nmole DNA oligo, 19 bases) were used for the amplification of the 16s rRNA sequence of the selected salt-tolerant bacterial isolates. The amplified products were purified and sequenced from Genelabs Medical, Sri Lanka by using Sanger Sequencing method. The Sanger Sequences obtained were analyzed and identified using BLAST search and were compared against bacterial 16s rRNA sequences available on NCBI database. The sequences were aligned by using Clustal W followed by construction of neighbor joining phylogenetic tree, using MEGA 11.

## Results

### Isolation of bacteria from soils of selected locations in the West coast of Sri Lanka

In the present study, a total of twelve soil samples from three locations; Negombo, Balapitiya and Beruwala on the West coast of Sri Lanka were screened for salt-tolerant bacteria (fig. 2). Soil samples were evaluated for their electrical conductivity (EC) and pH. The EC varied from 8.13 dS/m to 0.78 dS/m in soil samples collected from Negombo area. The EC varied from 1.44 dS/m to 0.47 dS/m in soil samples collected from Balapitiya whereas, soil samples from Beruwala area displayed an EC variation from 3.98 dS/m to 3.48 dS/m. The pH of different soil samples in three locations ranged from 7.74 to 7.36. (Table 3)

**Fig.2:**
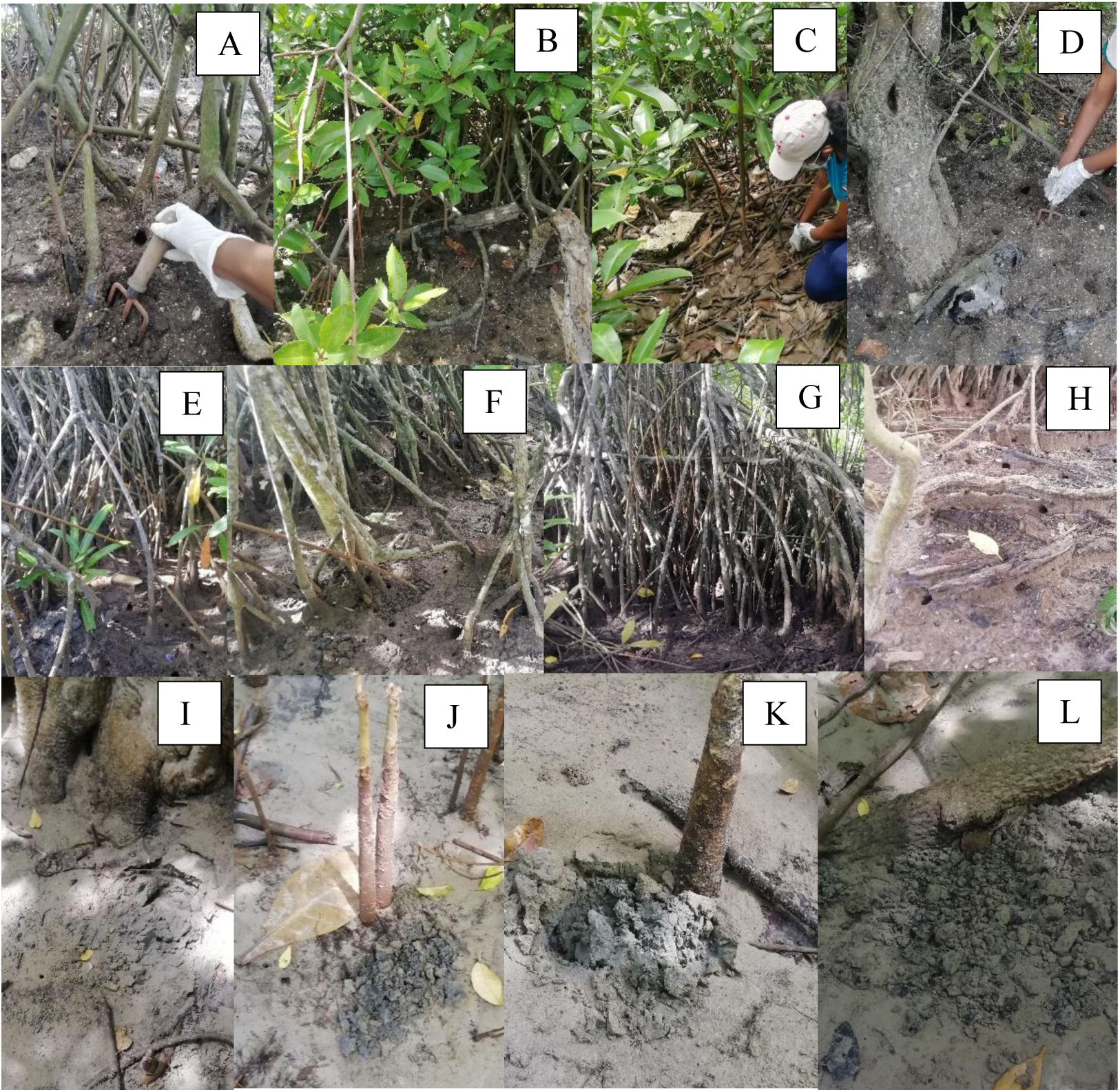
Soil sampling spots in three sampling locations. Note: spot 1 in Negombo (A), spot 2 in Negombo (B), s pot 3 in Negombo (C), spot 4 in Negombo (D), spot 1 in Balapitiya (E), spot 2 in Balapitiya (F), spot 3 in Balapitiya (G), spot 4 in Balapitiya (H), spot 1 in Beruwala (I), spot 2 in Beruwala (J), spot 3 in Beruwala (K) and spot 4 in Beruwala.

**Table 3:** Soil texture, pH and EC measurements and salinity classes of soils collected from three locations; Negombo, Balapitiya and Beruwala.

| Sampling spot No. | Location 01 – Negombo lagoon |  |  |  | Location 02 – Balapitiya |  |  |  | Location 03 – Beruwala |  |  |  |
| --- | --- | --- | --- | --- | --- | --- | --- | --- | --- | --- | --- | --- |
|  | Soil texture | Mean EC value (dS/m) | Mean pH value | Salinity class | Soil texture | Mean EC value (dS/m) | Mean pH value | Salinity class | Soil texture | Mean EC value (dS/m) | Mean pH value | Salinity class |
| 1 | Loamy with sands | 8.06 | 7.53 | Strongly saline | Loamy with sands | 1.44 | 7.62 | Non-saline | Sandy surface, loamy at 10 cm depth | 3.98 | 7.74 | Slightly saline |
| 2 | Loamy | 2.61 | 7.71 | Slightly saline | Loamy with sands | 1.31 | 7.57 | Non-saline | Sandy surface, loamy at 10 cm depth | 3.48 | 7.57 | Slightly saline |
| 3 | Clay | 0.78 | 7.56 | Non-saline | Loamy | 1.07 | 7.36 | Non-saline | Sandy surface, loamy at 10 cm depth | 3.79 | 7.74 | Slightly saline |
| 4 | Loamy with sands | 8.13 | 7.52 | Strongly saline | Sandy | 0.47 | 7.55 | Non-saline | Sandy surface, loamy at 10 cm depth | 3.52 | 7.72 | Slightly saline |

A total of 20 bacterial isolates were isolated from three locations. Maximum number of isolates were obtained from Negombo area (Table 4, Fig 3, Fig 4, Fig 5).

**Fig 3:**
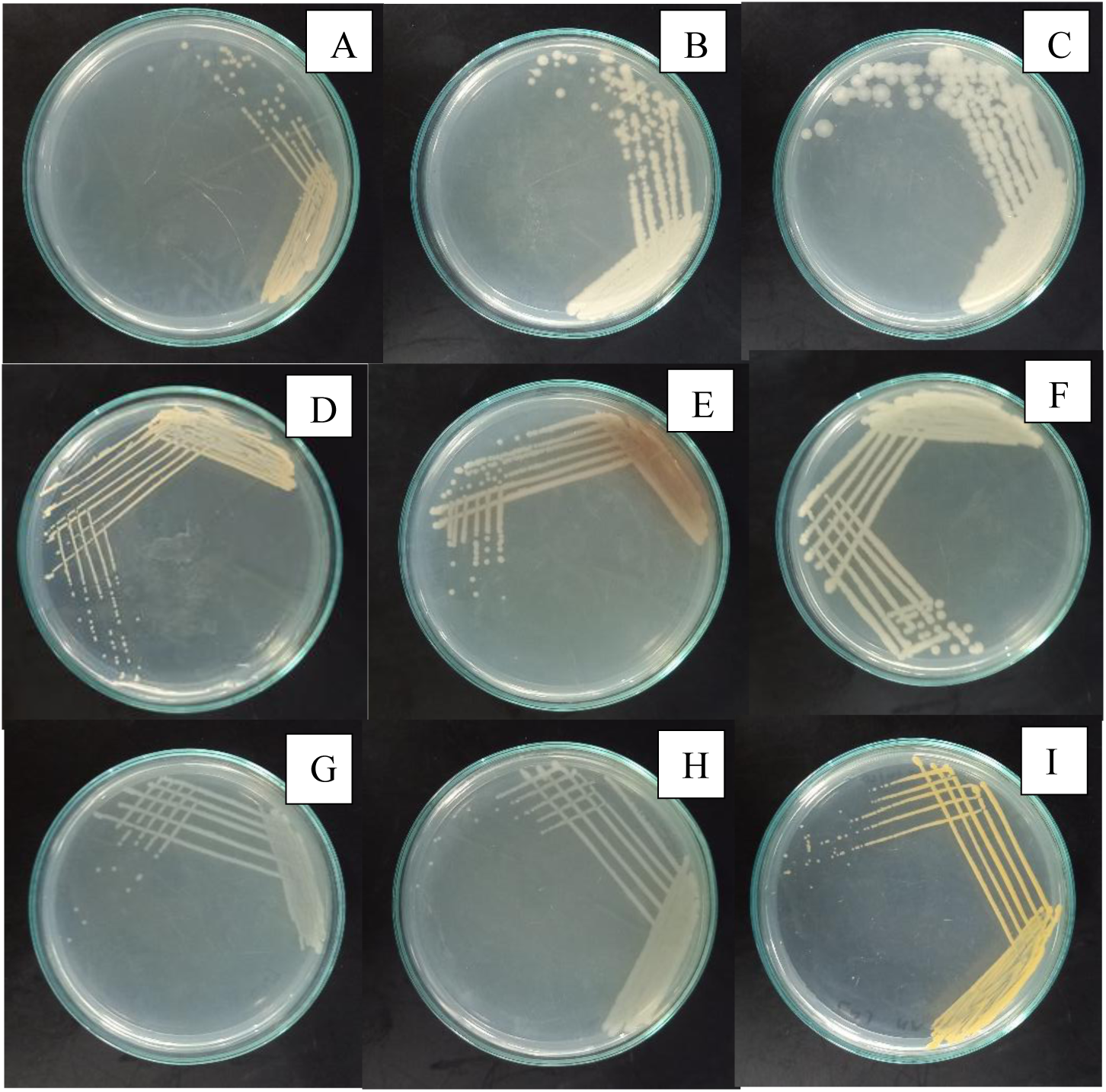
Bacterial isolates obtained from Negombo. Note: NE 01(A), NE 02(B),NE 03(C),NE 04(D),NE 05(E), NE 06(F), NE 07(G), NE 08(H), NE 09(I)

**Fig 4:**
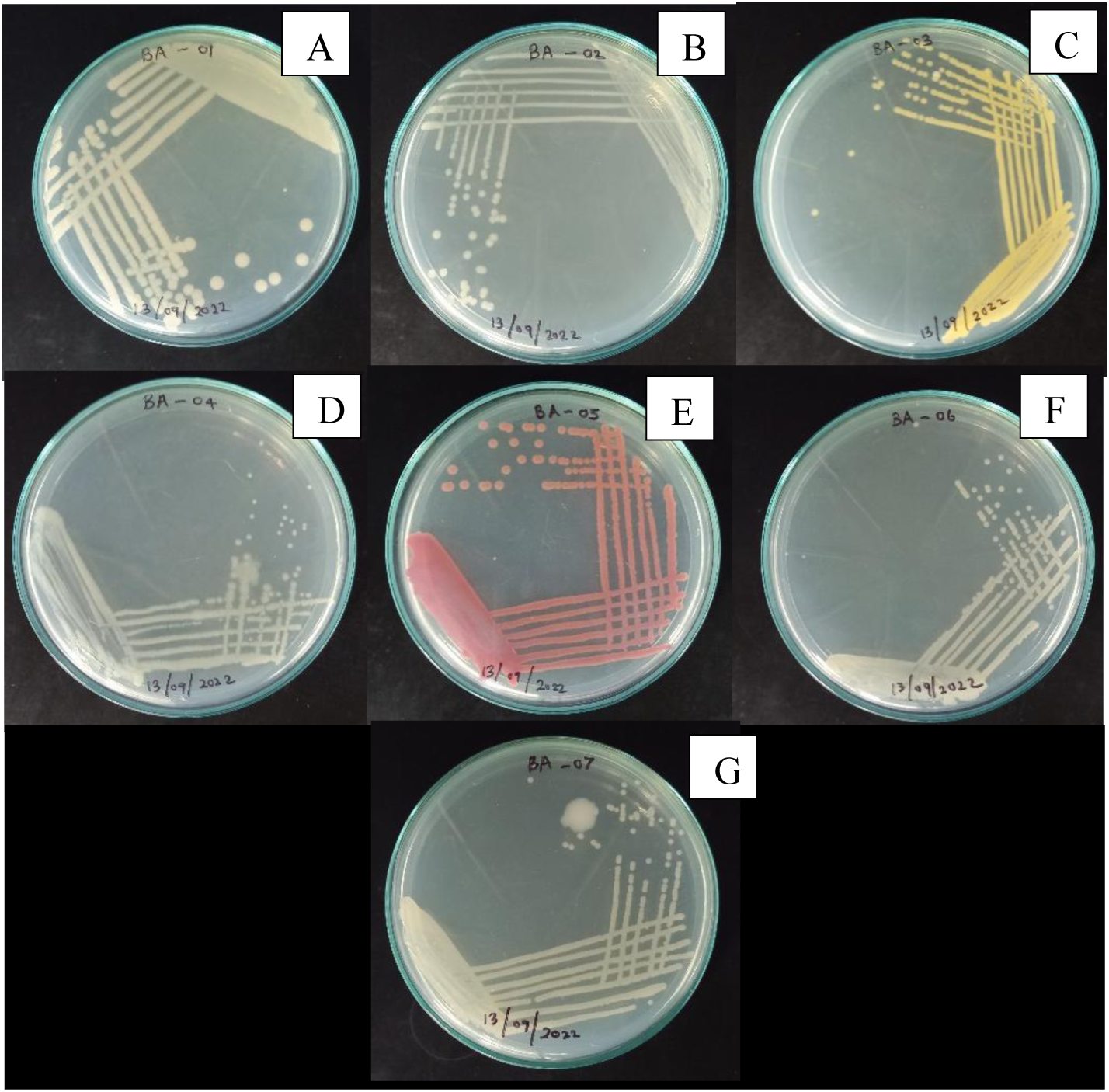
Bacterial isolates obtained from Balapitiya. Note: BA 01(A), BA 02(B), BA 03(C), BA 04(D), BA 05(E), BA 06(F), BA 07(G)

**Fig 5:**
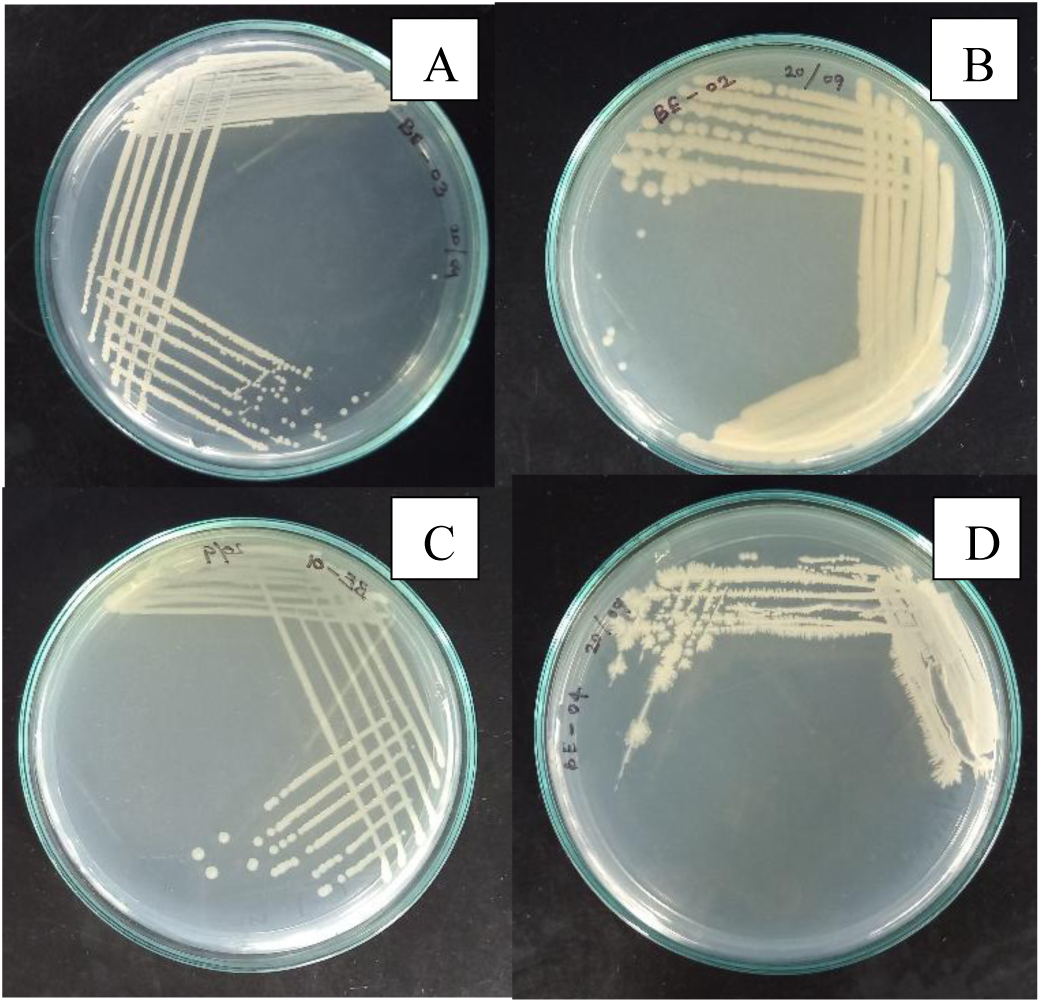
Bacterial isolates obtained from Beruwala. Note: BE 01(A), BE 02(B), BE 03(C), BE 04(D)

**Table 4:** Bacterial isolates obtained from 3 locations.

| Location | No. of isolates | Names of the isolates |
| --- | --- | --- |
| Negombo | 9 | NE 01, NE 02, NE 03, NE 04, NE 05, NE 06, NE 07 and NE 09 |
| Balapitiya | 7 | BA 01, BA 02, BA 03, BA 04, BA 05, BA 06 and BA 07 |
| Beruwala | 4 | BE 01, BE 02, BE 03 and BE 04 |

### Characterization of bacterial isolates

Macromorphological colony characters of the twenty isolates were observed and recorded (Table 5, Table 6, Table 7). Based on the Gram stain test, all the isolates were Gram-positive except BA 04. The twenty bacterial isolates were further characterized using biochemical tests namely, catalase test, modified oxidase test, hemolytic reaction on blood agar, and growth on MacConkey agar (Table 8, Table 9, Table 10).

**Table 5:** Macromorphological colony characters of bacterial isolates obtained from Negombo.

| Colony characteristics | Isolate name |  |  |  |  |  |  |  |  |
| --- | --- | --- | --- | --- | --- | --- | --- | --- | --- |
|  | NE 01 | NE 02 | NE 03 | NE 04 | NE 05 | NE 06 | NE 07 | NE 08 | NE 09 |
| Shape | Round | Round | Irregular | Round | Round | Round | Round | Round | Round |
| Size | Medium (1 mm) | Large (3 mm) | Large (3.5 mm) | Small (0.5 mm) | Medium (1 mm) | Large (4 mm) | Medium (1 mm) | Medium (1 mm) | Small (0.25 mm) |
| Margin | Entire | Undulating | Irregular | Entire | Entire | Entire | Irregular | Entire | Entire |
| Chromogenesis | Light orange | White | White | Light yellow | No pigmentation | Cream | No pigmentation | No pigmentation | Bright yellow |
| Opacity | Opaque | Opaque | Opaque | Opaque | Opaque | Opaque | Translucent | Translucent | Opaque |
| Elevation | Raised | Flat | Flat | Flat | Flat | Flat | Flat | Flat | Flat |
| Surface | Smooth | Dull | Dull | Smooth | Smooth | Smooth | Glistening | Smooth | Smooth |
| Consistency | Buttery | Buttery | Buttery | Buttery | Buttery | Buttery | Mucoid | Mucoid | Mucoid |

**Table 6:**
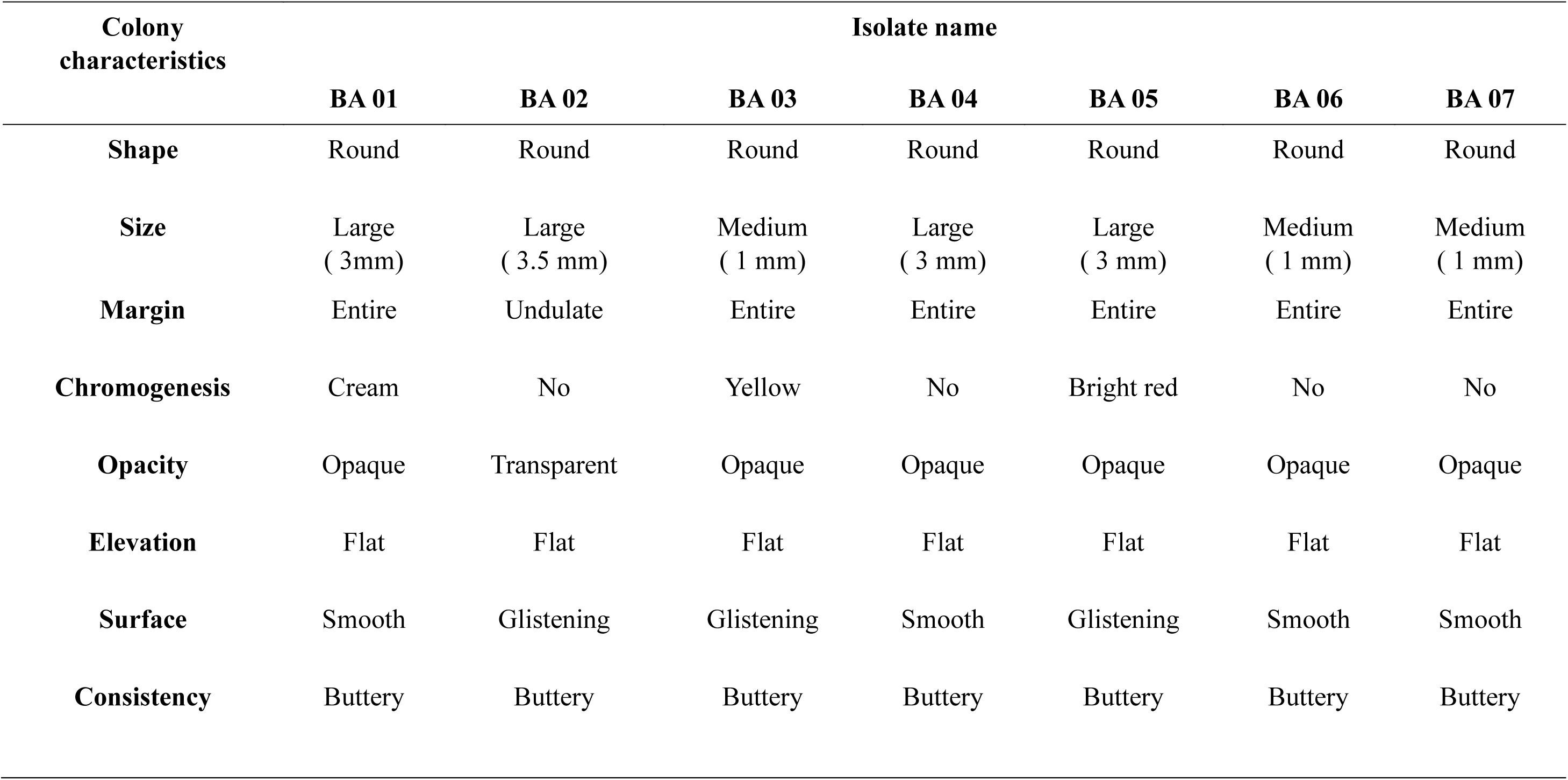
Macromorphological colony characters of bacterial isolates from Balapitiya.

**Table 7:** Macromorphological colony characters of bacterial isolates from Beruwala.

| Colony characteristics | Isolate name |  |  |  |
| --- | --- | --- | --- | --- |
|  | BE 01 | BE 02 | BE 03 | BE 04 |
| **Shape** | Round | Round | Round | Filamentous |
| **Size** | Small (0.5 mm) | Large (4 mm) | Medium (1 mm) | Large (3.5 mm) |
| **Margin** | Entire | Entire | Entire | Filamentous |
| **Chromogenesis** | White | Buff | No | White |
| **Opacity** | Opaque | Translucent | Transparent | Opaque |
| **Elevation** | Flat | Flat | Flat | Flat |
| **Surface** | Dull | Smooth | Smooth | Dull |
| **Consistency** | Brittle | Buttery | Mucoid | Brittle |

**Table 8:** Biochemical characterization of Gram positive cocci isolates.

| Isolate | Test |  |  |  |  |
| --- | --- | --- | --- | --- | --- |
|  | Gram's stain | Shape | Catalase | Modified oxidase | Hemolysis |
| NE 03 | Positive | Cocci | Negative | NA | $\gamma$ |
| NE 04 | Positive | Cocci | Positive | Negative | $\gamma$ |
| NE 06 | Positive | cocci | Negative | NA | $\alpha$ |
| NE 08 | Positive | cocci | Negative | NA | $\alpha$ |
| BA 05 | Positive | Cocci | Positive | Positive | $\alpha$ |
| BA 06 | Positive | Cocci | Negative | NA | $\gamma$ |
| BE 01 | Positive | cocci | Negative | NA | $\gamma$ |
| BE 02 | Positive | cocci | Positive | Positive | $\gamma$ |
| BE 03 | Positive | cocci | Negative | NA | $\alpha$ |
| BE 04 | Positive | cocci | Positive | Positive | $\alpha$ |

**Table 9:** Biochemical characterization of Gram positive bacilli isolates.

| Isolate | Test |  |  |
| --- | --- | --- | --- |
|  | Gram's stain | Shape | Hemolysis |
| NE 01 | Positive | bacilli | $\gamma$ |
| NE 02 | Positive | bacilli | $\alpha$ |
| NE 05 | Positive | bacilli | $\gamma$ |
| NE 07 | Positive | bacilli | $\gamma$ |
| NE 09 | Positive | bacilli | $\gamma$ |
| BA 01 | Positive | bacilli | $\alpha$ |
| BA 02 | Positive | bacilli | $\gamma$ |
| BA 03 | Positive | bacilli | $\gamma$ |
| BA 07 | Positive | bacilli | $\gamma$ |

**Table 10:** Biochemical characterization of Gram negative bacilli isolate.

| Isolate | Test |  |  |  |
| --- | --- | --- | --- | --- |
|  | Gram's stain | Shape | Hemolysis | Growth on MacConkey agar |
| BA 04 | Negative | bacilli | $\gamma$ | Non-fermenter |

### Screening of the isolates for salt tolerance

All twenty isolates were able to survive up to 20 dS/m (very strongly saline conditions) according to the absorbance values of the bacterial suspensions in different EC values taken at 600 nm (Table 11) and the growth of isolates under different EC values (Fig 6). According to the absorbance (600 nm) values, NE 03, NE 06, NE 07, BA 01 and BA 02 showed a significant increase of growth with increasing salinity and they were selected to observe the salinity tolerance behavior (Table 12). The highest increase in growth under very strongly saline conditions (20 dS/m) was observed in BA 01 followed by NE 06, BA 02, NE 07 and NE 03.

**Fig 6:**
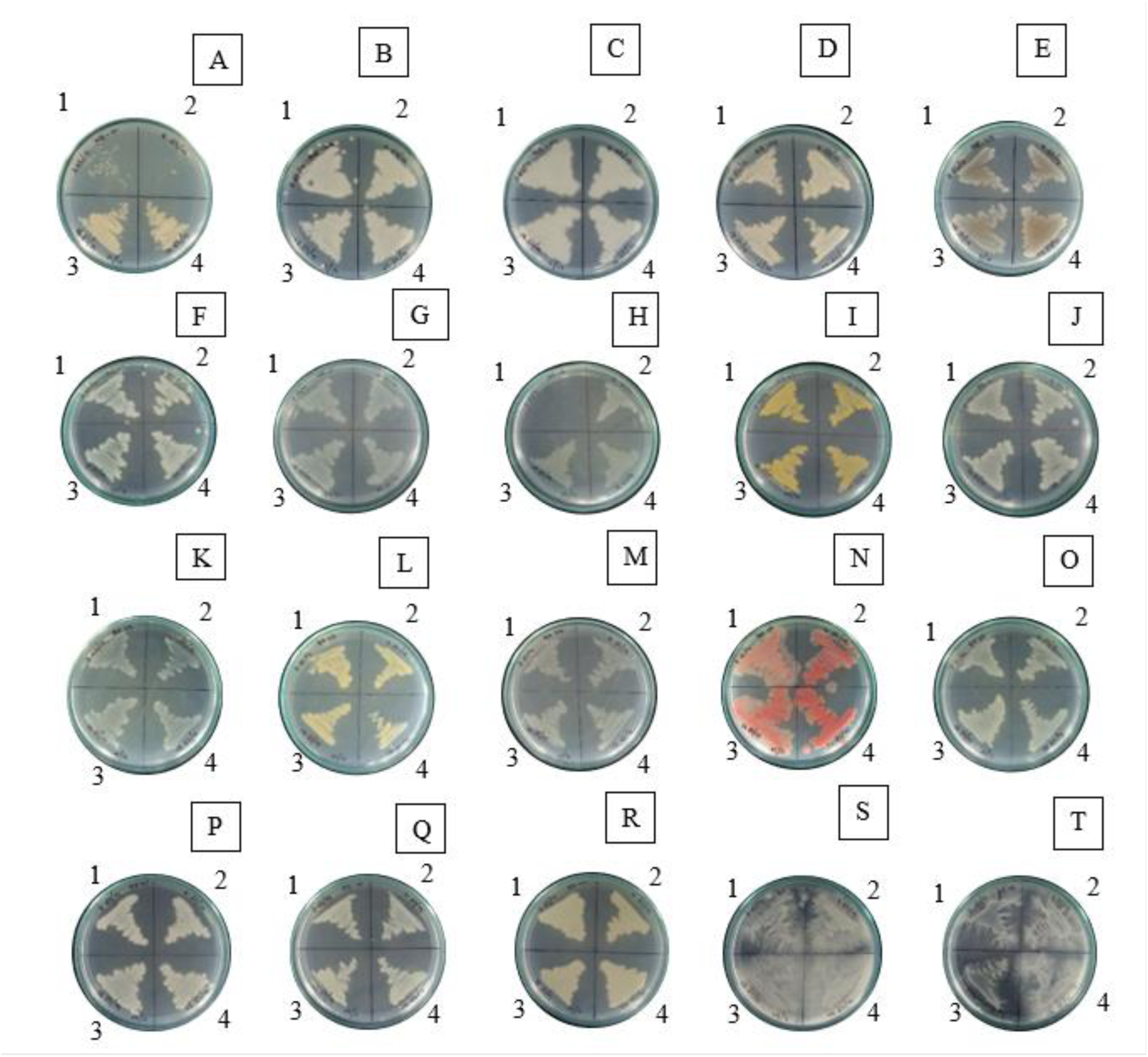
Growth of bacterial isolates under different concentrations of NaCl. Note: NE 01 (A),NE 02 (B),NE 03(C),NE 04 (D),NE 05 (E),NE 06 (F),NE 07(G),NE 08 (H),NE 09(I),BA 01(J),BA 02 (K),BA 03 (L),BA 04(M),BA 05 (N),BA 06 (O),BA 07(P),BE 01(Q),BE 02(R),BE 03(S),BE 04(T),3.13 dS/m(1), 7.16 dS/m(2), 13.98 dS/m(3), 20 dS/m(4)

**Table 11:**
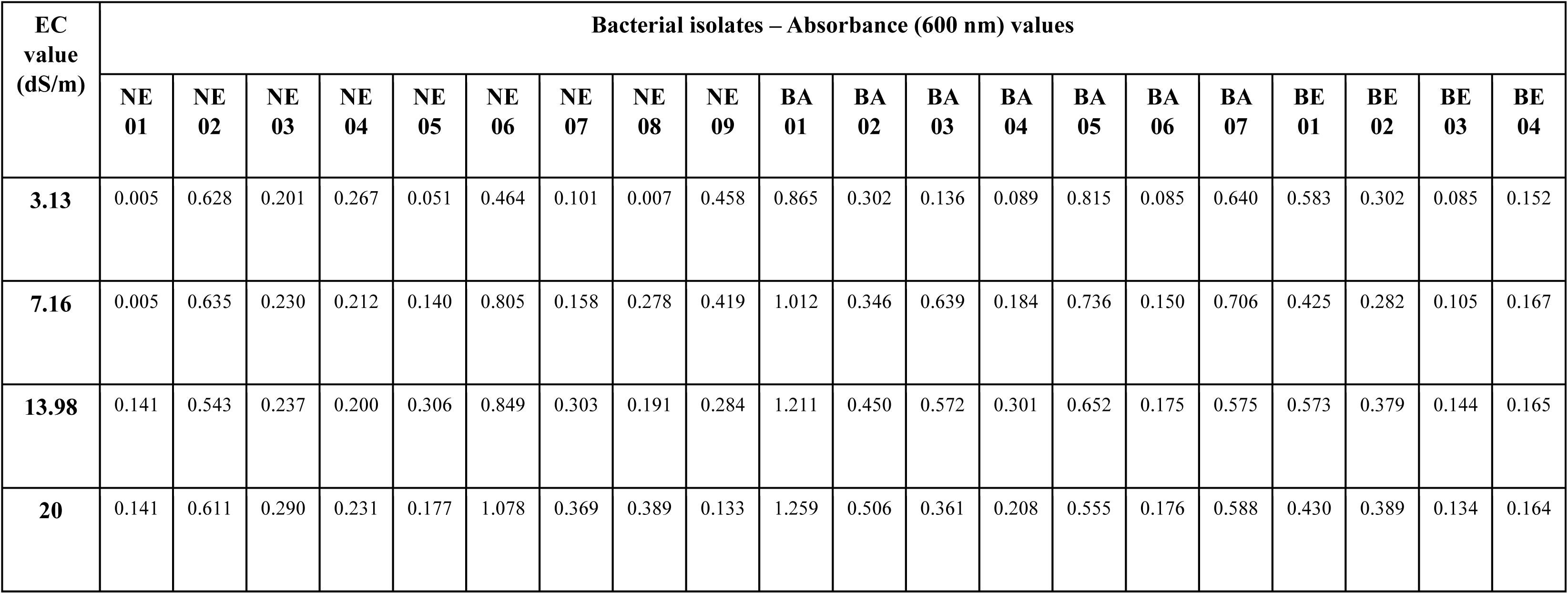
Absorbance (600 nm) values for 20 bacterial isolates under different EC values (salinity classes).

**Table 12:**
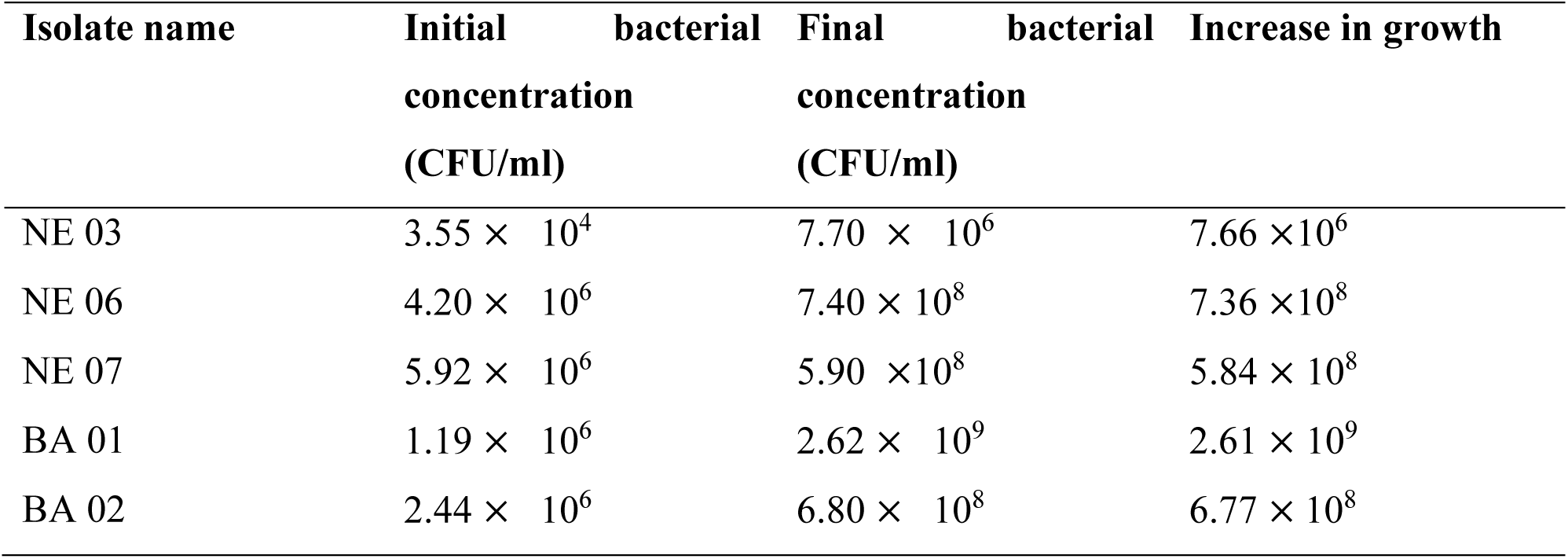
Salinity tolerance behavior of selected five salt tolerant isolates in (mention the salinity class).

### Molecular identification of selected isolates

Identification of the salt-tolerant isolates was based on Polymerase Chain Reaction (PCR) amplification of 16S rRNA gene sequences. The 16S rRNA gene of the selected isolates were successfully amplified using PCR, and approximately 1500 bp of the amplified products were sequenced (Table 13)). The BLAST-N comparison of the searched sequences in the NCBI nucleotide database revealed the isolate NE 03 showed 100% similarity with *Bacillus albus* strain BOE23 (NCBI accession number: OP919558.1), *Bacillus thurengiensis* strain VKK14 (NCBI accession number: OP906300.1) and *Bacillus proteolyticus* strain BP29 (NCBI accession number: OP905595.1). The isolate NE 06 showed 100% similarity with *Aeromonas* sp. strain 1051 (NCBI accession number: MT585851.1), *Aeromonas salmonicida* strain A1 (NCBI accession number: MT576565.1) and *Aeromonas sobria* strain A3-2 (NCBI accession number: MT576566.1). The isolate NE 07 showed 100% similarity with *Pseudomonas* sp. BL5 (NCBI accession number: LC547998.1), *Pseudomonas* sp. strain NIORKP101 (NCBI accession number: MH767183.1) and *Pseudomonas* sp. BMS12 (NCBI accession number: KU681078.1). The isolate BA 01 showed 99.69% similarity with *Aeromonas* strain NIO_15h (NCBI accession number: OP903270.1), *Aeromonas caviae* strain Mu-167 (NCBI accession number: OP889019.1) and *Aeromonas caviae* strain Mu-132 (NCBI accession number: OP889018.1). The isolate BA 02 showed 98.03% similarity with *Mangrovibacter yixingensis* strain MS 2.4 (NCBI accession number: KY321826.1), *Mangrovibacter* sp. MSSRF N97 (NCBI accession number: KU131267.1) and *Mangrovibacter plantisponsor* strain MSSFR N80 (NCBI accession number: KU131265.1). (Table 11). The phylogenetic tree of the selected bacterial isolates was constructed by using the neighbour-joining method (Fig. 8).

**Fig. 7:**
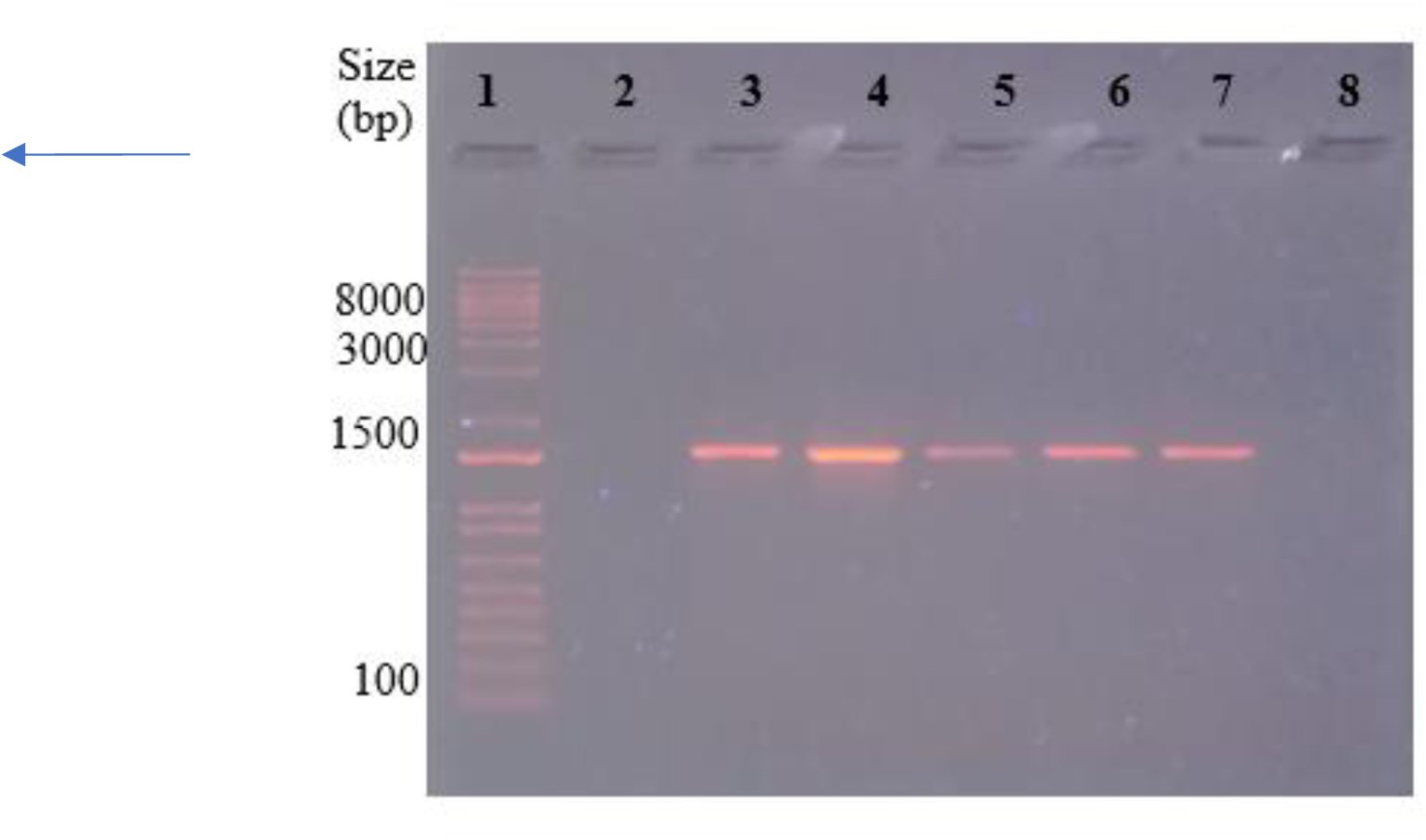
PCR amplified products of 16S rRNA of bacteria isolates in 1% agarose gel. 1kb DNA marker (1), blank (2), NE 03(3), NE 06(4), NE 07(5), BA 01(6), BA 02(7), negative control (8).

**Fig. 8:**
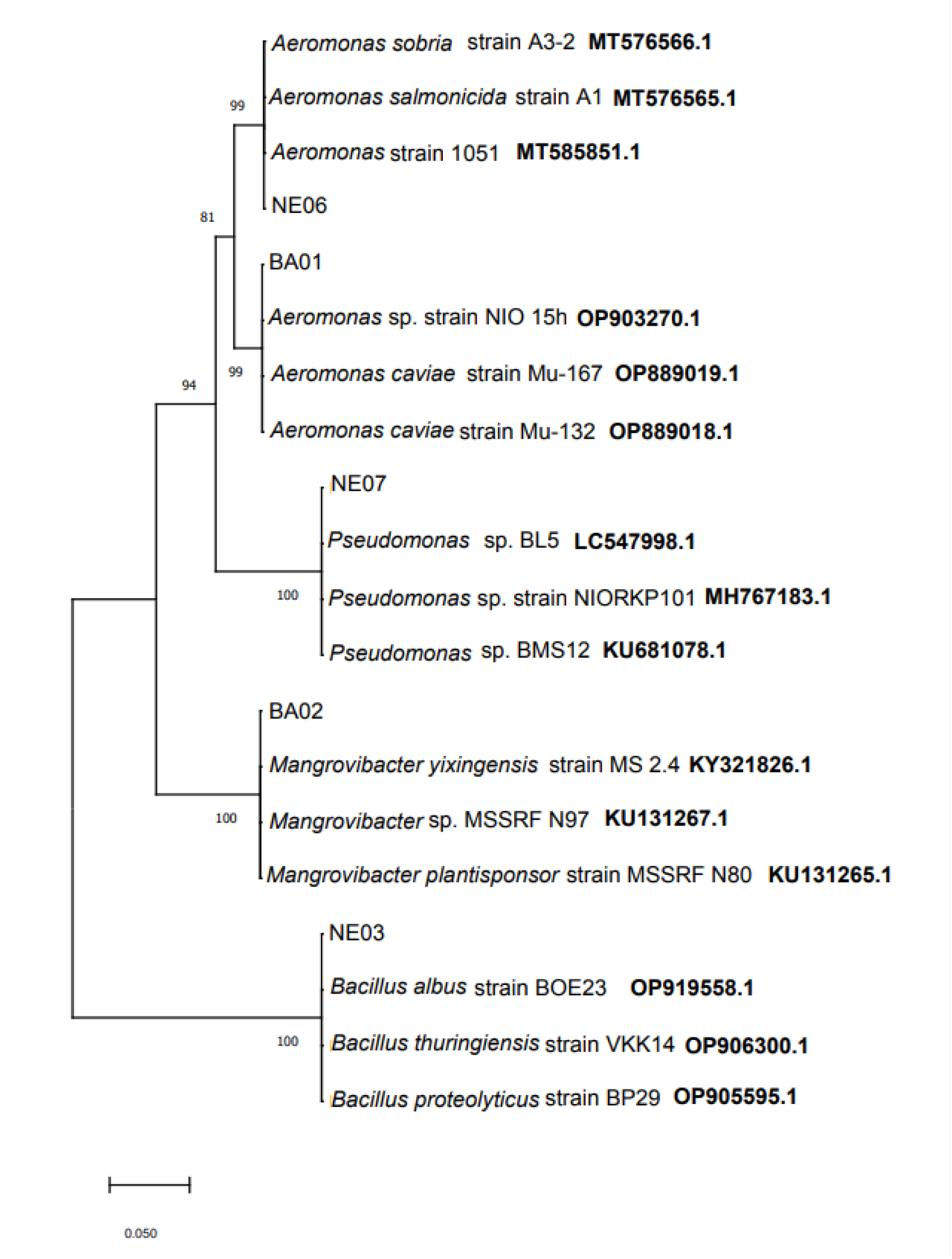
Phylogenetic tree constructed by the neighbour joining method derived from the analysis of the 16S rRNA gene sequence of salt tolerant bacterial isolates and related sequences obtained from NCBI. Scale bar 0.050, substitutions per nucleotide position.

**Table 13:** Identity profile of 16S rRNA gene partial sequences of selected five isolates according to the BLAST identification.

| Isolate Name | Sequence length(bp) | Bacterial species from GenBank database |  | Query cover | E value | Percentage identity |
| --- | --- | --- | --- | --- | --- | --- |
|  |  | Species name | Accession No |  |  |  |
| NE 03 | 1417 | *Bacillus albus* strain B0E23 | OP919558.1 | 100% | 1e-145 | 100.00% |
|  | 1464 | *Bacillus thurengiensis* strain VKK14 | OP906300.1 | 100% | 1e-145 | 100.00% |
|  | 1417 | *Bacillus proteolyticus* strain BP29 | OP90559.1 | 100% | 1e-145 | 100.00% |
| NE 06 | 1281 | *Aeromonas* sp. Strain 1051 | MT585851.1 | 100% | 1e-165 | 100.00% |
|  | 1420 | *Aeromonas salmonicida* strain | MT576565.1 | 100% | 1e-165 | 100.00% |
|  | 1401 | *Aeromonas sobria* strain A3-2 | MT576566.1 | 100% | 1e-165 | 100.00% |
| NE 07 | 1415 | *Pseudomonas* sp. BL5 | LC547998.1 | 100% | 9e-157 | 100.00% |
|  | 699 | *Pseudomonas* sp. strain NIORKP101 | MH767183.1 | 100% | 9e-157 | 100.00% |
|  | 1525 | *Pseudomonas* sp.BM512 | KU681078.1 | 100% | 9e-157 | 100.00% |
| BA 01 | 1385 | *Aeromonas* strain NIO_15h | OP903270.1 | 100% | 1e-165 | 99.69% |
|  | 1406 | *Aeromonas caviae* strain Mu-167 | OP889019.1 | 100% | 1e-165 | 99.69% |
|  | 1408 | *Aeromonas carviae* strain Mu-132 | OP889018.1 | 100% | 1e-165 | 99.69% |
| BA 02 | 863 | *Mangrovibacter yixingensis* | KY321826.1 | 100% | 6e-149 | 98.03% |
|  | 1387 | *Mangrovibacter* sp. MSSRF N97 | KU131267.1 | 100% | 6e-149 | 98.03% |
|  | 1360 | *Mangrovibacter plantisponsor* strain MSSRF N80 | KU131265.1 | 100% | 6e-149 | 98.03% |

## Discussion

In the present study, three locations in the West coast of Sri Lanka; Negombo, Balapitiya and Beruwala were selected as representative locations. As the West coast of Sri Lanka is mainly divided into North West coast and South West coast, Negombo was selected from North West coast, Balapitiya was selected from South West coast and Beruwala was selected as a location between North West coast and South West coast. The soil sampling in Negombo was carried out during Southwest monsoon season; the month of September. The sampling location was a narrow mangrove area close to the Negombo lagoon. Soils from sampling spots 1 and 4 can be categorized into strongly saline class, soils in sampling spot 2 is under slightly saline class and soils in sampling spot 3 is under non-saline class. Therefore, there is a considerable salinity variation among soil sampling spots at the same location. The reason for this salinity variation could be the influence of freshwater runoff from the land during monsoons to the mangrove area. The soil collected from sampling spot 3 displayed a clay texture. Clay (soil) can hold more fresh water (include reference) which can be the reason for lower salinity levels compared to the other three sampling spots.

In the present study, all twenty isolates from three locations were able to survive upto the highest saline concentration tested (20 dS/m). From the selected five isolates, the highest increase in growth under 20 dS/m were observed in BA 01, followed by NE 06, BA 02, NE 07 and NE 03.

In a similar study; (Sharma et al., 2021), isolated, screened and characterized salt tolerant bacteria with plant growth promoting activities from saline agricultural fields of Haryana, India. A total of thirteen isolates were found salt tolerant at different salt concentrations out of 81 bacterial isolates. In their study, the isolates which has significant salt tolerant and PGPR ability were identified as *Bacillus paramycoides*, *Bacillus amyloliquefaciens* and *Bacillus pumilus*. Leontidou et al., 2020, reported that bacterial strains of *Sarcocornia* sp, *Atriplex* sp. And *Crithmum* sp. can tolerate salinity ranged from 65 mS cm^-1^ to 72 mS cm^-1^. Do et al., 2022, found that the diazotropic salt tolerant bacterial strains of *Klebsiella, Agrobacterium, Pseudomonas* and *Ochrobacterium* isolates from the roots of a halophytic plant. *Arthrochemum indicum* showed salinity tolerance ranging from 4% to 8% NaCl the productivity of peanut in saline and control conditions has improved (Pal et al., 2021). Egamberdieva et al., 2019 explored that, the genetic diversity of salt tolerant plant growth promoting rhizobacteria (ST-PGPR) and found that most of the isolates were able to tolerate up to 8% NaCl and belong to the genus *Bacillus.* Zhang et al., 2018 reported that 162 bacterial strains out of total 305 bacterial strains were tested for salt tolerant up to 150g/l NaCl concentration. Further the researchers conducted a phylogenic analysis of 74 of these salt tolerant strains and showed that they belong to orders *Bacillales, Actinimycetales, Rhizobiales and Oceanospirillales*. Most of the isolates showed their potential in improving salt tolerance, growth and yield of rice under salt stress conditions.

Similarly, several researchers reported the diversity of ST-PGPR in the coastal areas. For example, in Andaman and Nikobar islands of India, isolated 23 bacterial strains, out of total 121 bacterial strains showed salt tolerance up to 10% NaCl with plant growth promoting characteristics including production of indole acetic acid (IAA), siderophore, extracellular enzymes and phosphate solubilization and it revealed that the majority of isolates were *Bacillus* sp. (Amaresan et al., 2016) However, in one study, all the isolated bacteria have shown salt tolerant up to 1.2% salt concentration which belongs to very strongly saline category (> 16 dS/m), (Wang et al., 2021)

Although several researchers tested bacterial strains for salt tolerant in combination with plant growth promoting activities, we did not perform salt tolerant studies with plant growth promoting activities.

Soil salinity has a serious impact on plant growth and the uptake of high amount of salt inhibits physiological and metabolic processes of plants. However, there are several conventional methods of reclamation of saline soils such as flushing, leaching, or adding an amendment (gypsum, CaCl2) are successful to some extent as well as adversely affect the agro-ecosystems (Egamberdieva et al., 2019). In the present study, the bacterial isolates were further screened for their salt tolerance to salts at different salt concentrations ranging from 3 dS/m to 20 dS/m. The optical density (at 600 nm) of the bacterial suspensions in different concentrations of NaCl measured at the end of the five-day incubation period. The absorbance values indicate the rate of bacterial growth at different salinity concentrations. The salinity tolerance behavior with increasing salinity based on absorbance values at 600 nm for 20 bacterial isolates were graphically illustrated.

The isolates; NE 03, NE 06, NE 07, BA 01 and BA 02 with significant salt tolerant ability were sequenced for amplified 16S rRNA gene and identified upto the genus level as, *Bacillus sp.*, *Aeromonas sp., Pseudomonas sp., Aeromonas sp. and Mangrovibacter sp.* respectively.

The growth of bacteria in the media containing high salt concentrations is an important indicator of their salt tolerance. According to relevant studies, the biomass of bacteria will change as they adapt to new osmotic environments (Weinisch et al., 2018). Further, some halophilic bacteria change their morphology to adapt to the high salinity conditions as cell lengthening has been observed in *Pseudomonas sp.(Manter, 2009; Li et al., 2014). Mccready et al. (1966)*. Pumping out ions from the cell is another strategy for avoiding high salt concentrations in the cytoplasm (Hunte et al., 2005). Some microorganisms produce organic osmotic substances that accumulate in the cytoplasm and resist osmotic pressure under high salt stress such as betaine (Wu et al., 2018) and proline (Nadeem et al., 2007). A study conducted by Jiang et al., 2010 found that *Bacillus sp.* accumulate proline under high salt concentrations. Therefore, for the present study, an analysis of the accumulation of betaine and proline in NE 03, NE 06, NE 07, BA 01 and BA 02 will prove that they can thrive in saline water.

In a similar study conducted by Das et al., 2015, seven isolates (D-TSB-84, D-TSB-86,H-TSB-70,P-TSB-70,P-TSB-72,P-TSB-75 and P-TSB-78) were found extreme (8-20%) NaCl stress tolerant. Out of seven salt tolerant bacteria, three isolates (P-TSB-70, P-TSB-72 and P-TSB-75) tolerated 20% NaCl stress and four isolates (D-TSB-84, P-TSB-78,D-TSB-86 and H-TSB-70) tolerated 9% NaCl. According to 16S rRNA sequence results, the seven extreme salt bacteria were documents into genus *Staphylococcus, Enterococcus, Enterobacter* and *Proteus*. Further, the study revealed that, two isolates belong to genus *Staphylococcus* and one isolate belong to *Proteus* grew well in 20% salt stressed medium. However, these workers did not perform screening of bacteria isolates for their PGPR potential. They focused on the marine environment to obtain diverse microbial strains with variable stress tolerant properties and they concluded that these identifies salt stress tolerant bacteria would be highly useful in obtaining salt stress tolerant genes in future to develop the salt stress tolerant transgenic plant varieties. In another study conducted by Amaresan et al., 2016 isolated a total of 121 rhizobacteria from tsunami affected Andaman and Nikobar islands. Among 121 isolates, 91 isolates were able to tolerate 5% NaCl and 23 isolates were able to tolerate 10% NaCl. The 23 isolates were further screened for the PGP properties; phosphate solubilization, production of IAA, siderophore production and extracellular enzymes production. The researchers used 16S rDNA sequencing to identify the efficient salt tolerant bacteria belonged to *Bacillus* spp. Followed by *Alcaligenes* sp., *Microbacterium* sp. and *Lysinibacillus* sp. Further, they performed nif H region amplification of salt tolerant PGPR to confirm the harboring gene responsible for atmospheric nitrogen fixation. Out of 23 salt tolerant isolates only 3 isolates which belong to *Bacillus sp.* and *Enterobacter sp.* showed positive results. On the basis of PGP properties, three isolates showed positive to all PGP traits which all of them belong to *Bacillus sp.* Therefore, The Bacillus sp. proved potential salt tolerant PGP rhizobacteria. Since PGPR have value in wide range of applications by inducing plants to resist saline environmental conditions, in future studies these isolates can be screened for their PGP activities thereby the positive isolates will use in the process of development of a biofertilizer.

(And these isolates can be used in the process of development of bioferlizers and can be used in saline soil for growth promotion of agricultural crops).

### Conclusion

In this study, 20 soil bacterial isolates were isolated from the selected locations in the West coast of Sri Lanka namely Negombo, Balapitiya and Beruwala. All the 20 bacterial isolates were found as salt tolerant up to the highest NaCl concentration tested which was 18 dS/m. Among these 20 bacterial isolates only five selected isolates namely NE 03, NE 06, NE 07, BA01 and BA 02 were tested increase in growth at 18 dS/m and found that the highest increase in growth was in BA 01 followed by NE 06, BA 02, NE 07 and NE 03. The selected five salt tolerant isolates were identified with their closely related species using sequence analysis followed by BLAST-N comparison. According to the results obtained from BLAST-N comparison, it could be able to identify the genus of the selected five isolates correctly. Therefore, NE 03 belongs to the genus Bacillus, both NE 06 and BA 01 belong to genus Aeromonas, NE 07 belongs to genus Pseudomonas and BA 02 belongs to genus Mangrovibacter. By considering the identified genus of selected five isolates there is a potential of having PGPR activities in both Bacillus and Pseudomonas species. Therefore, in future studies, the identified Bacillus and Pseudomonas bacterial strains can be further screen for their PGPR activities and can be used to formulate biofertilizers to boost crop productivity in salt affected saline agricultural lands.

## Notes

### Competing Interest Statement

The authors have declared no competing interest.

### Funding

- John Keells Research, Nanotechnology and Science Park, Mahenwatta, Pitipana, Homagama, Sri Lanka

