## Supplementary Material for "Isolation and Characterization of Salt Tolerant Soil Bacteria from Selected Locations in the West Coast of Sri Lanka"

### APPENDIX 01

#### 1.1 Media preparation

1.1.1 **Preparation of nutrient agar**

A weight of 28.0 g of nutrient agar powder were suspended in 1000 ml of distilled water. The solution was heated to boiling to dissolve the medium completely. Then it was sterilized by autoclaving at 15 lbs pressure, 121 °C for 20 minutes. The solution was cooled to 45-50 °C, mixed well and poured into sterile petri plates.

#### 1.1.2 Preparation of blood agar

A weight of 28 g of nutrient agar powder were suspended in 1000 ml of distilled water. The solution was heated to boiling to dissolve the medium completely. It was sterilized by autoclaving at 15 lbs pressure, 121 °C for 20 minutes. The nutrient agar medium was cooled to 45-50 °C and 5% (V/V) sterile defibrinated blood was added that has been warmed to room temperature and the mixture was mixed well. The solution was dispensed into sterile petri plates.

#### 1.1.3 Preparation of MacConkey agar

A weight of 50 grams of MacConkey agar powder was suspended in 1000 ml of distilled water. The solution was heated to boiling to dissolve the medium completely. It was sterilized by autoclaving at 15 lbs pressure, 121 °C for 20 minutes. The solution was cooled to 45-50 °C, mixed well and poured into sterile petri plates.

#### 1.1.4 Preparation of LB broth without NaCl solution

Requirement of ingredients to prepare 500 ml of LB broth without NaCl as follows; Yeast extract = 2.5 g

Tryptone = 5.0 g

The above measured ingredients were suspended in 500 ml of distilled water. The solution was heated to boiling to dissolve the medium completely. It was sterilized by autoclaving at 15 lbs pressure, 121 °C for 20 minutes. The solution was cooled to room temperature (25 ± 2°C).

#### 1.1.5 Preparation of LB broth solutions with different concentrations of NaCl

Final EC values of LB solutions were decided based on the table below.

#### Table 1.1.5 Final EC value of LB for four solutions based on salinity classes.

| **Solution No.** | **Saturate extract (dS/m)** | **Salinity class** | **Decided final EC value of**  **LB (dS/m)** |
| --- | --- | --- | --- |
| 1 | 2-4 | Slightly saline | (2+4)/2 = 3 |
| 2 | 4-8 | Moderately saline | (4+8)/2= 6 |
| 3 | 8-16 | Strongly saline | (8+16)/2=12 |
| 4 | > 16 | Very strongly saline | (16+20)/2=18 |

To calculate the NaCl concentration for a decided EC value, Total Dissolved Solids (TDS) in mg/l = K * EC

If EC < 5 dS/m, K= 640 If EC > 5 dS/m, K = 800

EC of LB without NaCl = 3 dS/m

The above value was taken as a reference value and NaCl concentration in each solution was calculated as follows;

For solution 1,

LB broth was taken without adding NaCl For solution 2,

TDS = K * EC

= 800 * (6-3) dS/m

= 2400 mg/l

Required concentration of NaCl = 2.4 g/l For solution 3,

TDS = K * EC

= 800 * (12-3) dS/m

= 7200 mg/l

Required concentration of NaCl = 7.2 g/l For solution 4,

TDS = K * EC

= 800 * (18-3) dS/m

= 12,000 mg/l

Required concentration of NaCl = 12 g/l

#### 1.1.6 Preparation of LB for a serial dilution to take CFU/ml

Requirement of ingredients to prepare 500 ml of LB, Yeast extract = 2.5 g

Tryptone = 5 g NaCl = 5 g

The above measured ingredients were suspended in 500 ml of distilled water. The solution was heated to boiling to dissolve the medium completely. It was sterilized by autoclaving at 15 lbs pressure, 121 °C for 20 minutes. The solution was cooled to room temperature (25 ± 2°C).

#### 1.2 Preparation of serial dilutions

1.2.1 **Preparation of a serial dilution to culture the bacteria from soil samples** A serial dilution was prepared by using sterile distilled water upto 10^-3^.

Three test tubes each containing 9 ml of sterile distilled water were prepared and labelled as 10^-1^,10^-2^ and 10^-3^. The volume of 1 ml from original soil suspension was added to 10^-1^ test tube. From 10^-1^ test tube, 1 ml volume was added to 10^-2^ test tube. This process was continued upto 10^-3^.

#### 1.2.2 Preparation of a serial dilution to obtain CFU/ml

Nine test tubes each containing 9 ml of sterile LB were prepared and labelled as 10^-^ ^1^,10^-2^,10^-3^ upto 10^-9^. The volume of 1 ml from original bacterial suspension was added to 10^-1^ test tube. From 10^-1^ test tube, 1 ml volume was added to 10^-2^ test tube. This process was continued upto 10^-9^.

### APPENDIX 02

#### 2.1 Measuring Electrical Conductivity (EC) of soil samples.

2.1.1 **Instrument used**

Digital EC meter (Hanna instruments HI199301)

#### 2.1.2 Main features of the instrument

Accuracy: 2%

Measurement range: 0 to 20 mS/cm

Resolution: 0.1 mS/cm

Battery life: approximately 250 hours

#### 2.1.3 Preparation of soil solutions to measure EC

The weight of 10 g soil was added into 10 ml of distilled water and a soil solution was prepared for each sample. Solutions were mixed well by vortexing.

#### 2.1.4 Taking measurements by using EC meter

The probe of the EC meter was washed with distilled water and blot dried with dust free tissues/kimwipes prior to use. Then the probe was dipped into the soil solution and the EC value was taken for a particular soil solution after displaying “stable” on digital screen.

#### 2.2 Measuring pH of soil samples

2.2.1 **Instrument used**

pH meter

2.2.2 **Main features of the instrument** Accuracy pH mV Â°C: ± 0.01 pH ± 1 mV Temperature Compensation: 0 to 100°C Input Impedance: > 10 ohms

#### 2.2.4 Preparation of soil solutions to measure pH.

A weight of 10 g soil was added into 10 ml of distilled water and a soil solution was prepared for each sample. Solutions were mixed well by vortexing.

#### 2.2.5 Taking measurements by using pH meter.

After calibrating the pH meter, both the pH probe and temperature probe were rinsed with distilled water and blot dried with dust free tissues/kimwipes. Two probes were dipped into a particular soil solution and the pH value was taken after displaying a stable value on the digital screen.

#### 2.3 Measuring absorbance at 600 nm using UV- visible spectrophotometer

2.3.1 **Main features of the instrument**

Optical system: Double beam (1200 lines/mm grating) Wavelength range: 190- 1100 nm

Absorbance accuracy: ± 0.005 (for absorbance values below 1.0 A)

#### 2.3.2 Taking absorbance values at 600 nm by using UV- visible spectrophotometer

First the instrument was programmed to measure the absorbance at 600 nm. Then a blank cuvette which is containing NaCl without LB broth solution was placed in the spectrophotometer. The option of “0 ABS” was clicked to read 0.00000 A. The sample which is containing bacteria was poured into another cuvette and it was placed in the spectrophotometer. After that, the absorbance value can be taken for the particular sample.

### APPENDIX 03

#### 3.1 Bacteria culturing techniques

3.1.1. **Spread plate technique**

A dilution series were prepared from a sample. The volume of 0.1 ml from the appropriate desired dilution was pipetted out onto the surface of an agar plate. The sample was spread evenly over the surface of agar using the sterile glass spreader while rotating the petri dish carefully. The plate was incubated at 30 °C for 24 hours.

#### 3.1.2 Four quadrant streak plate technique

A loop was flamed until red hot and let to cool. A single colony was taken into the loop and streaked as at A in figure 2.1.2. The loop was re-flamed and let it cool. The plate was streaked as at B to spread the original inoculum over more of the agar. Again, the loop was re-flamed and let it cool. Streaking was performed as at C. The loop was re- flamed and let it cool. The plate was streaked as at D. (figure 2.1.2) The plate was inverted and incubated at 30 ° C for 24 hours.

**APPENDIX 04**

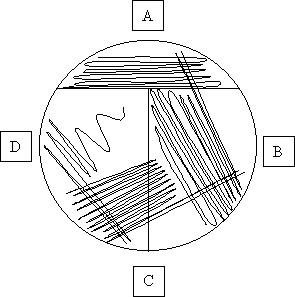

**Figure 3.1.2: Four quadrant streak plate**

### APPENDIX 04

**Table 4: Colony morphology of bacterial isolates**

| **Bacterial colony**  **characteristics** | **Description** |
| --- | --- |
| Colony shape | Includes the form, elevation and margin  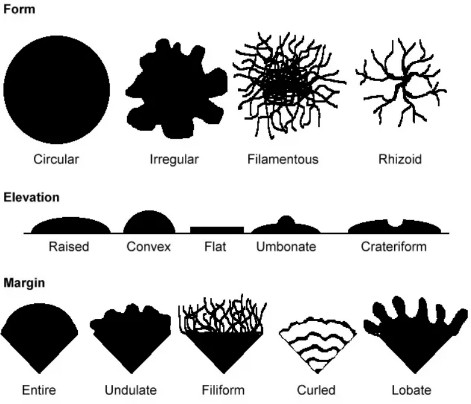 |
| Size of the bacterial colony | The diameter(d) of a representative colony d < 1mm – small  d = 1 mm – medium  d > 1 mm – large |
| Chromogenesis | Color of the colonies (pigmentation) |
| The opacity of the bacterial colony | Transparent(clear)  Opaque (not transparent or clear)  Translucent (almost clear, but distorted vision) Iridescent (changing colors in reflected light) |
| Surface of the colony | Dull (opposite of glistening) Veined  Rough  Wrinkled (or shriveled) Glistening |
| Consistency of the colony | Dry Moist  Viscid (sticks to loop, hard to get off)  Brittle/friable (dry, breaks apart) Mucoid (sticky, mucus-like) |

### APPENDIX 05

#### 5.1 Gram stain test

5.1.1 **Reagents used in gram staining**
      - Crystal violet; the primary stain
      - Gram’s iodine; the mordent
      - A decolorizer made of acetone and alcohol (95%)
      - Safranin/corbol fucision (dilute); the counter stain

#### 5.1.2 Procedure

A clean, grease free slide was taken. A thin smear of suspension was prepared with a loopful of bacterial sample. The slide was air dried, and heat fixed. Crystal violet was poured onto the slide and kept for about 1 minute and rinsed with water. Then the grams iodine was flooded, kept for about 1 minute and washed with the water. Acetone was added to the slide and rinsed with water after keeping about 10-20 seconds. The counter stain was added, the slide was kept about 1 minute and rinsed with water. The slide was air dried, blot dried and observed under light microscope (total magnification: 1000X)

#### 5.2 Catalase test

5.2.1 **Reagents used in catalase test**
      - 3% H2O2

#### 5.2.2 Procedure

Slide method

A sterile wooden stick was used to transfer a small amount of colony growth into the surface of a clean, dry glass slide. Then a drop of 3% H2O2 was placed in the slide. Evolution of oxygen bubbles was observed.

#### 5.3 Modified oxidase test

5.3.1 **Reagents used in modified oxidase test**
      - Filter	paper	disks	impregnated	with	tetramethyl-p-phenylenediamine dihydrochloride (oxidase reagent)

#### 5.3.2 Procedure

A filter paper disk was transferred into a petri dish by using a sterile forcep. A small amount of several colonies was rubbed on the filter paper disk. The filter paper disk was incubated at room temperature for 2 minutes. The color development was observed.

### APPENDIX 06

#### 6.1 Reconstitution of Proteinase K

Proteinase K powder was resuspended in Molecular Biology Grade Water to obtain a 20 mg/ml stock solution.

To find the required amounts for 5 preparations,

Required amount of Proteinase K for 20 preparations = 10 mg

Required amount of Proteinase K for 5 preparations = (10 mg/20) *5 = 2.5 mg

Required volume of molecular biology grade water for 20 preparations = 0.5 ml Required volume of molecular biology grade water

for 5 preparations	= (0.5 ml/20) * 5 = 0.125 ml

#### 6.2 Preparation of Lysozyme solution [for gram positive bacteria only]

The volume of 1 ml Lysozyme solution was prepared by dissolving 45 mg of lysozyme powder (MB098) in 1ml of gram positive lysis solution . The mixture was pipetted up and down to dissolve the lysozyme.

#### 6.3 Preparation of diluted wash solution

To find the required amounts for 5 preparations,

Required volume of wash solution for 20 preparations = 3 ml

Required volume of wash solution for 5 preparations = (3 ml/20) * 5 = 0.75 ml Required volume of 100% ethanol to dilute wash solution for 20 preparations,

=12 ml Required volume of 100% ethanol to dilute wash solution for 20 preparations,

= (12 ml/20) * 5

= 3 ml

#### 6.4 Gram positive bacterial preparation

The volume of 1.5 ml of bacterial broth culture was pelleted in provided 2 ml capped collection tube by centrifuging for 2 minutes at 13,000 rpm at room temperature (15-25 °C). The pellet was resuspended thoroughly in 200 μl of lysozyme solution and it was incubated for 30 minutes at 37 °C. The volume of 20 μl Proteinase K solution was added to the sample. Then 20 μl of RNase A solution was added, mixed and incubated for 5 minutes at room temperature (15-25 ° C). A volume of 200 μl lysis solution was added and the sample was mixed by vortexing for few seconds. It was incubated at 55 °C for 10 minutes. Then the DNA isolation procedure was followed.

#### 6.5 Gram negative bacterial preparation

The volume of 1.5 ml of bacterial broth culture was pelleted in provided 2 ml capped collection tube by centrifuging for 2 minutes at 13,000 rpm at room temperature (15-25 °C). The pellet was resuspended thoroughly in 180 μl of lysis solution (AL)

. The volume of 20 μl proteinase K solution was added to the sample, mixed and incubated for 30 minutes at 55 °C. The volume of 20 μl RNase A solution was added to the sample, mixed and incubated for 5 minutes at room temperature (15-25 ° C). The volume of 200 μl lysis solution (C1) was added to the sample, it was mixed thoroughly by vortexing about 15 seconds and incubated at 55 °C for 10 minutes. Then the DNA isolation procedure was followed.

#### 6.6 DNA isolation from gram positive and gram negative bacteria

The volume of 200 μl of ethanol (95-100%) was added to the lysate and mixed thoroughly by vortexing for few seconds. The lysate was transferred onto HiElute Miniprep Spin column (capped) provided. The lysate was centrifuged at 10,000 rpm for 1 minute at room temperature (15-25 °C). The flow through liquid was discarded and the spin column was placed in same 2 ml collection tube. The volume of 500 μl prewash solution was added to the column, and it was centrifuged at 10,000 rpm for 1 minute at room temperature (15- 25 °C). The flow through liquid was

discarded and the same collection tube with the column was re-used. The volume of 500 μl diluted Wash Solution (WS) was added to the column and it was centrifuged for 3 minutes at 13,000 rpm at room temperature (15-25 °C). (If the solution has not completely passed through the membrane, the sample should be centrifuged again at higher speed until all the solution has passed through).The flow through liquid was discarded and the sample was centrifuged again at same speed for additional 1 minute to dry the column. Then the HiElute Miniprep Spin Column (capped) was transferred to fresh collection tube. The volume of 200 μl elusion buffer was added directly into the column without spilling to the sides. It was incubated for 1 minute at room temperature (15 -25 °C). The column containing collection tube was centrifuged at 10,000 rpm for 1 minute at room temperature (15-25 °C) to elute the DNA. The elute was transferred to a fresh capped 2 ml collection tube for longer use. The tube containing elute was stored at -20 °C.

#### 6.7 DNA quantification using fluorometer.

First fluorometer must be calibrated by using “Blank” followed by “Standard DNA (100 ng/μl)” sample provided.

Requirement of solutions for samples as follows;

For “Blank” sample; the volume of 100 μl dye and 100 μl of TE buffer should be added. Total volume should be 200 μl.

For “Standard DNA” sample; the volume of 100 μl dye, 98μl of TE buffer and 2 μl from “standard DNA” should be added. Total volume should be 200 μl.

For each DNA sample which needs to be quantified; the volume of 100 μl dye, 98μl of TE buffer and 2 μl from DNA sample should be added. Total volume should be 200 μl.

#### 6.7.1 Preparation of TE buffer

Concentration of stock solution = 20 X Concentration of working solution = 1X

Assumption: required volume of working solution = 10 ml

C1V1	= C2V2

20X * V1 = 1X * 10 ml V1	= 0.5 ml

According to the above calculation, 0.5 ml of TE buffer was required from the concentrated solution in order to prepare 10 ml of working solution. The volume of

0.5 ml TE buffer was added into sterile falcon tube and the remaining 9.5 ml volume was filled with sterile distilled water.

#### 6.7.2 Preparation of dye

The dye should be freshly prepared before adding it to the samples.

Concentration of stock solution = 200 X Concentration of working solution = 1X

Assumption: Two DNA samples need to quantify

Therefore, Volume of dye should be sufficient for four samples including “Blank” and “Standard” samples.

Total volume of dye required for four samples = (100 μl * 4) = 400 μl

C1V1	= C2V2

200X * V1 = 1X * 400 μl

V1	= 2 μl

According to the above calculation, 2 μl was required from stock solution in order to prepare 400 μl of working solution. TE buffer was required to dilute the dye. Therefore, TE buffer requirement was (400- 2) μl = 398 μl.

After preparing the working solution containing the dye, the tube was covered with aluminum foils to avoid degradation.

After preparing the working solutions, the required volumes from each solution were added to the blank, Standard and DNA samples which need to be quantified. Since the dye was inside each tube, all the tubes should be covered with aluminum foils. First “Blank” sample was inserted into the fluorometer for calibration. Then “standard DNA” sample was inserted into the fluorometer for calibration. After that, bacterial genomic DNA samples were quantified by using the fluorometer.
